# Continuous visualization of differences between biological conditions in single-cell data

**DOI:** 10.1101/337485

**Authors:** Tyler J. Burns, Garry P. Nolan, Nikolay Samusik

**Affiliations:** Department of Cancer Biology, Stanford University School of Medicine, Stanford CA, USA; Department of Microbiology and Immunology, Stanford University School of Medicine, Stanford CA, USA

## Abstract

In high-dimensional single cell data, comparing changes in functional markers between conditions is typically done across manual or algorithm-derived partitions based on population-defining markers. Visualizations of these partitions is commonly done on low-dimensional embeddings (eg. t-SNE), colored by per-partition changes. Here, we provide an analysis and visualization tool that performs these comparisons across overlapping k-nearest neighbor (KNN) groupings. This allows one to color low-dimensional embeddings by marker changes without hard boundaries imposed by partitioning. We devised an objective optimization of k based on minimizing functional marker KNN imputation error. Proof-of-concept work visualized the exact location of an IL-7 responsive subset in a B cell developmental trajectory on a t-SNE map independent of clustering. Per-condition cell frequency analysis revealed that KNN is sensitive to detecting artifacts due to marker shift, and therefore can also be valuable in a quality control pipeline. Overall, we found that KNN groupings lead to useful multiple condition visualizations and efficiently extract a large amount of information from mass cytometry data. Our software is publicly available through the Bioconductor package Sconify.

## Introduction

Emerging technologies are providing high parameter information from single cells, providing many opportunities to study the diversity of complex biological systems. These technologies include mass cytometry (1), high-parameter imaging systems (2-4), and single-cell sequencing (5).

A growing body of dimension reduction and manifold embedding algorithms have been adapted to these methods such as force-directed graphs (6) (7), t-SNE (8,9), and principal component analysis (PCA) (10) to represent the distribution of cells in high-dimensional space near each other in two dimensions. These methods provide a relatively simple visualization of an otherwise complicated data type.

To perform statistical comparisons across multiple samples in this type of data, researchers often first partition the concatenated dataset into disjoint subsets (clusters or gates) based on surface markers, and then for each subset performing sample-to-sample comparison of markers that were not used for the partitioning (functional markers, often phospho-proteins) (11,12). This family of approaches allows for statistical analysis of discrete changes such as signaling differences between subsets or differential abundance of cell subsets(13-19) (20) (21) (22) (23).

The results of clustering can be visualized directly on single-cell manifold embeddings, where each cell is colored by information within the cluster it belongs to. However, clustering and gating is subject to ambiguity depending on the choice of a gating strategy or clustering algorithm. Furthermore, a visualization of change across conditions on a low-dimensional embedding contains hard boundaries defined by the partitions. An alternative approach is the use of overlapping cell-centered neighborhoods rather than disjoint clusters, allows for a comparison with potentially higher granularity, which doesn’t depend on a cell’s membership in a specific group. One example of this is Cydar, a method that tests for differential abundance of cells across biological conditions using per-cell overlapping hyperspheres in high dimensional data(24).

A related and intuitive approach to analyzing a high-dimensional dataset is by grouping each data point by its respective nearest neighbors in feature space, which can in turn be used for comparisons. The concept of nearest neighbor-based analysis, particularly for classification, was first described in a treatise authored by Hasan Ibn al-Haytham (Alhazen) roughly one millennium ago during the Islamic Golden Age (25). Despite its medieval roots, it is still competitive with modern and more complex analysis tools (26,27).

Here, we present a computational method for single cell data that groups cells by their KNN and focuses on changes in marker expression across biological conditions. This method specifically produces t-SNE maps that one can color by KNN-based changes across conditions. Thus, our tool is geared at visual enhancement of lower dimensional embeddings.

Our method is called SCONE, which stands for **S**mooth **C**omparison **o**f **Ne**ighbors, with our Bioconductor package accordingly called “Sconify.” SCONE makes statistical comparisons (e.g. signaling differences in basal versus perturbed cells) in overlapping KNN around single cells. Thus, each cell represents the information contained within itself and its local neighborhood. The method concatenaties multiple biological conditions, and performs KNN using population-defining markers (e.g. surface markers) as input that are not expected to change between conditions. Functional marker comparisons are then performed for each KNN.

These per-cell KNN changes are then concatenated to the expression matrix used as input. The user has the choice to run t-SNE on this expression matrix as well. While the software provides internal visualization capabilities using the ggplot2 package, the output is a csv file that can be uploaded into visualization software like Cytobank, FlowJo, or CYT. SCONE adds to a growing list of single-cell analysis methods which employ nearest-neighbor computation, including density-based clustering (7), smoothing (28) and network-based community detection (29).

This work serves not only to introduce a visual tool, but to explore the uses, benefits, and limitations of these KNN-based comparisons applied directly to CyTOF data with respect to functional marker responses and quality control based on differential cell abundance. While these aspects are being addressed separately by an increasing number of algorithms in computational flow cytometry (30), the value of KNN applied to CyTOF data is that one can have access to this information along with aforementioned information (e.g. density) within a single method. Therefore, our method coupled to a lower dimensional visualization like t-SNE is especially useful in the early stages of data analysis, where the intent is often to quickly and efficiently gain intuition about the data quality, subset diversity, and the scale of differences across comparisons of interest.

We re-analyzed a pre-existing B cell development dataset to highlight a small subset highly responsive to IL-7 through pSTAT5 (31). We also re-analyzed a PBMC dataset created while CyTOF was still relatively new, utilizing KNN as an evaluation metric for data quality and normalization efficacy (32). We synthetically altered this dataset to assess the sensitivity of KNN as a data quality metric. Overall, we show that relatively simple and intuitive nearest neighbor-based analysis can be effectively utilized in mass cytometry for both quality control and marker expression comparisons.

## Methods

### Mass cytometry experiments

Mass cytometry data used in this manuscript, together with the information regarding cell preparation, data acquisition, and processing was obtained from the original publications (31-33). Through the rest of the manuscript, we will call these respective datasets by their main contributing authors: Bodenmiller/Zunder/Finck, Bendall/Davis/Amir, and Fragiadakis.

### Data input

A schematic of the algorithm’s workflow is provided (Figure 1A). In brief, cells from a basal condition and one or more stimulatory conditions are concatenated into a single matrix of cells by features, with an additional column denoting condition, using the FlowCore Bioconductor package. The software produces this matrix from a list of fcs files the user provides as input. These cells are then subject to an appropriate normalizing transformation (e.g. arsinh-transformed with a cofactor of five, which became a *de facto* standard for CyTOF analysis(1)). The user then has the option to do per-marker quantile normalization (34) and/or z-score transformation, as a means to correct batch effects and reduce sample-to-sample variability (see **Correcting for technical artifacts in the data** for efficacy analysis). Each cell’s k-nearest neighbors are determined with user-chosen input markers (in our case, surface markers) using Euclidean distance in the R implementation Fast Library for Approximate Nearest Neighbors (flann) R package. Importantly, the our software uses an exact KD-tree based KNN finder within the rflann package, not an approximate KNN finder.

**Figure 1:**
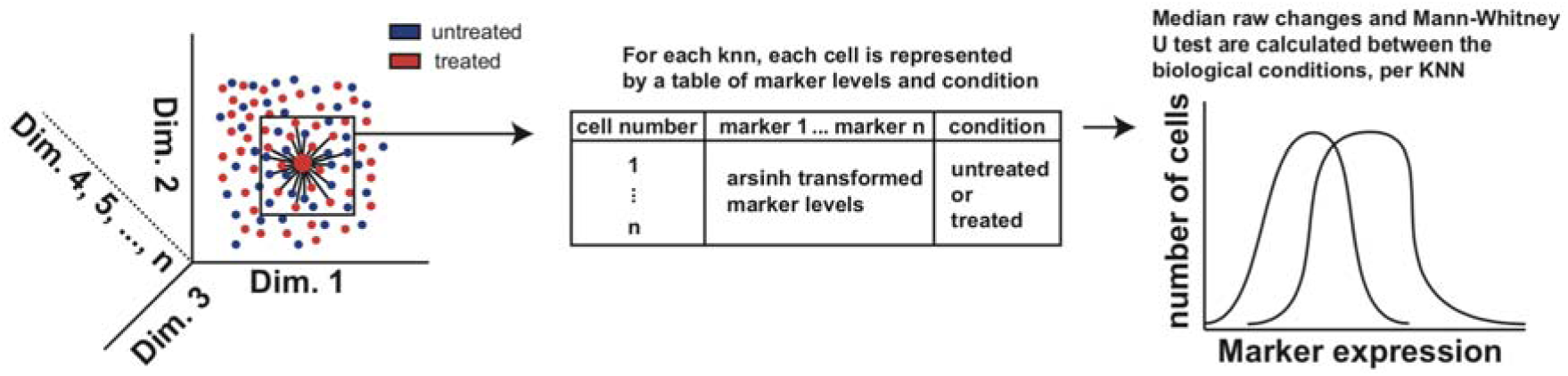
Schematic of the SCONE algorithm and its output. (left) Cells from two or more biological conditions are used as input for k-nearest neighbors (KNN) generation, using user-defined features. (middle) Each KNN is a matrix of cells by features, which include functional markers hypothesized to change between conditions, along with an additional column demarcating which biological condition was used. (right) Statistical tests are performed between the distributions per feature, and Arsinh differences are thresholded by respective FDR-adjusted q values.

### Paired sample comparison of functional marker distributions per KNN

For each k-nearest neighborhood, for each functional marker of interest, two values are calculated. The first is the raw change between the two sample, which is defined as the difference between the median values of two samples, with one sample typically representing the baseline condition and the other some sort of perturbation (e.g. a stimulation)..” The second value is the p-value output from a user-chosen statistical test (currently, Mann-Whitney U test (35) or T test (36)) between the distributions of marker values for each biological condition. Accordingly, the p-value is adjusted for false discovery rate using p.adjust function within the R stats package. We therefore call the statistical test output q-values. Following this, we give the user the option to threshold the raw changes by q-value. In other words, if the q-value for a given k-nearest neighborhood is lower than a user-defined cutoff (e.g. 0.05), then the raw change will be reported. Otherwise, the raw change will take on the value zero.

### Per-donor comparisons

In experimental setups with multiple donors or replicates, the user has an option to make a per-sample (rather than per-cell) comparison between two conditions. For each k-nearest neighborhood, for each marker of interest, the user-chosen median or mean values of expression are computed for each sample designated as “control” condition and each sample designated as an “experimental” condition (e.g. stimulated cells). These values are then used as input in a t-test comparing control versus experimental per-sample expression for the given marker. The resulting output is the per-sample comparison p-values. Like the paired sample p-values, they are corrected for false discovery rate using the Benjamini-Hochberg methods within the p.adjust function in the R stats package. They are therefore also referred to as q-values, as shown in basal versus LPS-treated healthy human whole blood (33) (Figure S1).

### Structure of SCONE output

The results of comparisons (both median change and q-values) are appended column by column to the end of the original single cell expression matrix (the data matrix of the FCS file). Each new column is either a q-value or a raw change of a single functional marker and therefore can be parsed in the same way as other cytometry data. The software also provides a density estimation (1 divided by average distance to KNN){Samusik:2016ev}, and a readout of differential abundance of cells belonging to one condition versus the other. The user then has the option to run t-SNE within our software using a wrapper for the Rtsne R package, which automatically adds the t-SNE embedding coordinates as columns to the end of the data matrix. Of note, the user has the option to subsample the data prior to the t-SNE step as well, in order to minimize runtime(30). This can be done independent of the KNN-based analysis as needed. We recommend the user choose the same input markers for t-SNE as those that went into the KNN calculation. Although t-SNE is an effective way to reduce high dimensional data into two dimensions, there are other methods that could just as effectively visualize the data that are beyond the scope of this manuscript, and other methods may be more optimal than t-SNE depending on the biological question and the number of cells being used as input (37). We intend to update the software package with additional dimension reduction methods as they become relevant to the mass cytometry field.

After SCONE runs to completion, these data can then be analyzed directly in R through, for example, various analysis and visualization tools developed and/or collected in the Bioconductor CyTOF workflow (21). These data can also be exported as a csv and used in external visualization tools like FlowJo, Cytobank (DROP) (38), or the MATLAB-based CYT (9,29,31).

### Selection of the number of nearest neighbors

We chose the optimal number of nearest neighbors (k) by solving the functional marker imputation problem, i.e. by determining how well the functional response markers could be predicted from the respective values of the k-nearest neighbors. Specifically, for each cell we computed mean values of the functional variables from its k-nearest neighborhood. We then optimized the *k* value (neighborhood size), using the Euclidean distance between the actual functional variable vector for a given cell and the imputed median vector as a loss function (39) (Figure S2A).

We evaluated this optimization on a previously published mass cytometry dataset consisting of human PBMCs across a variety of stimulatory conditions on the Bodenmiller/Zunder/Finck and Fragidakis datasets. We observed that across a range of *k* values ranging from n/5 to n/1000, the relationship between log10(k) and the imputation loss was parabolic, with a clearly defined minimum. As such, one could find the value of k that effectively minimized the imputation loss (Figure S2B). We found that for n = 10,000 PBMCs in these data, the ideal k was determined to range between n/20 to n/100 depending on stimulatory condition.

We expect this value to differ depending on datasets and biological conditions being used. If the calculated ideal k value differs between biological conditions being compared, we recommend choosing a value k that is the mean of ideal k values found across the biological conditions. Within our software, we provide the user the ability to use this metric for the dataset in question prior to executing the KNN statistics, and we recommend this be done with each new dataset.

### Using KNN to assess and correct technical artifacts

There may be shifts in marker expression levels between biological conditions, that result entirely from technical artifacts. This could potentially affect the validity of the statistical comparisons within each cell’s KNN. However, we found that our KNN framework can be pre-emptively used to identify such artifacts, and determine the efficacy of any corrections performed accordingly, as a means of quality control and assurance of the validity of the downstream analysis.

We assess the presence of technical artifact due to marker shifting using the per-KNN fraction of cells belonging to each non-baseline biological condition compared to the user-designated baseline for each KNN. If we have two files belonging to the same donor, perhaps using a short-term stimulatory condition where surface marker expression is not expected to change, then we can hypothesize that for each KNN in the concatenated data, roughly half of the cells should belong to one or the other condition. As a control, we subsample a single data file into two matrices without replacement, and pretend they are two separate data files. If we run KNN (n = 10,000 cells, k = 100) on these data and plot a histogram of the per-KNN percent of cells belonging to “condition 2,” we find that the standard deviation (0.05) is what would be expected by the binomial distribution with k independent trials and probability of 0.5, under the De Moivre-Laplace theorem (Figure 2A) (40). Thus, we use this standard deviation as our definition of “best possible” manifold overlap. The following section describes in technical detail how this is used to calculate our manifold overlap score (m score) for data quality analysis. Having established those means to assess the manifold overlap, we also provide the user with the ability to perform per-marker quantile normalization, and/or z-score transformation of each feature going into the KNN generation. The Sconify package can perform these transformations on the data independent of the rest of the pipeline as needed. We see this as an important pre-processing and quality assurance step primarily with datasets in which multiple donors are combined.

**Figure 2:**
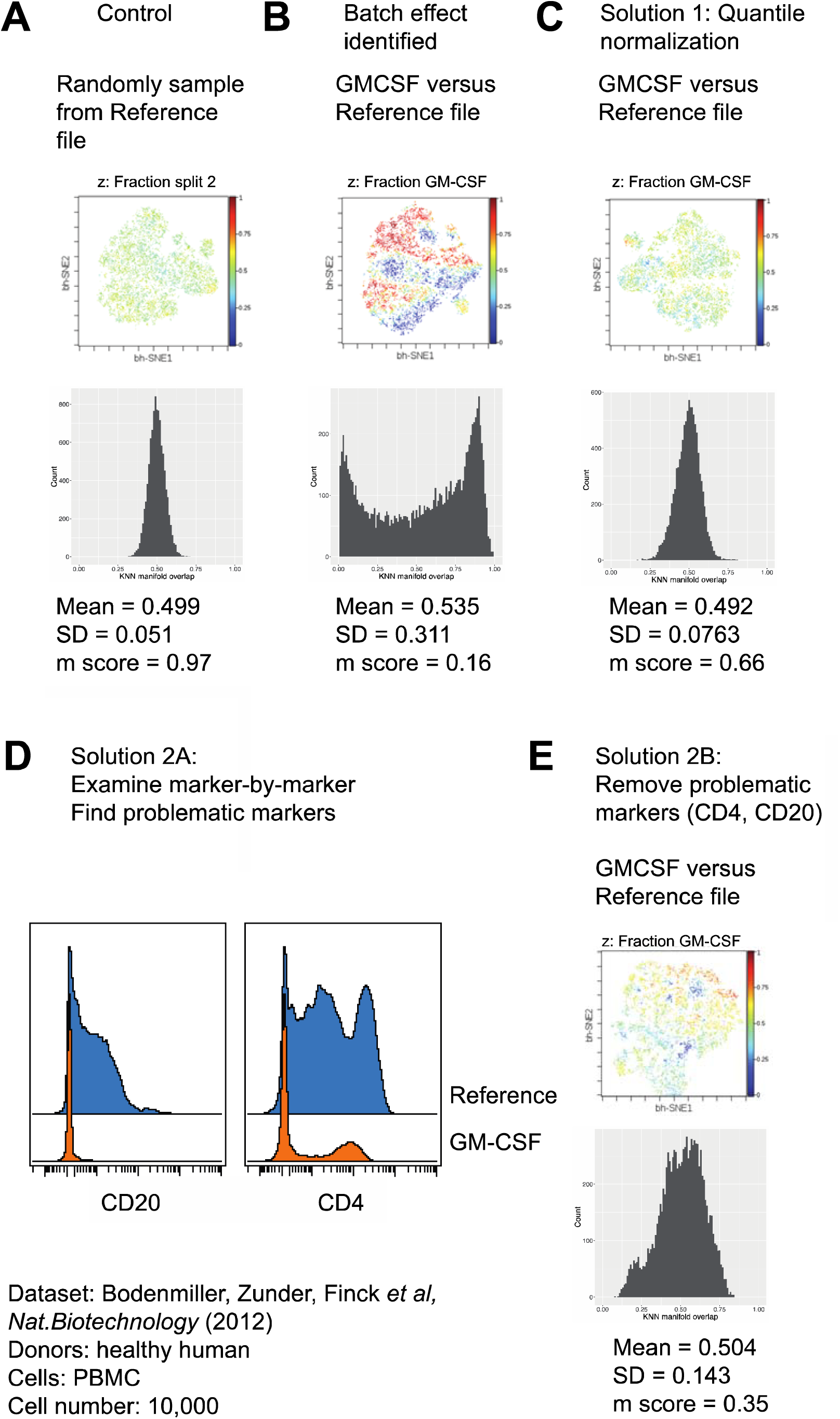
A KNN (k = 100) metric for manifold overlap. (A) Split-file control: a single file subsampled into two separate matrices without replacement. Differential abundance of cells belonging to one matrix or the other is assessed with KNN, and plotted as a histogram along with a t-SNE map colored by KNN manifold overlap. (B) Untreated versus GM-CSF treatment assessed in the same manner reveals a distribution with two peaks and a substantially higher standard deviation (and therefore lower *m* score). (C) These same files run again after per-marker quantile normalization of surface markers reveal an improved *m* score. (D) Per-marker inspection reveals alterations between files in CD4 and CD20. These markers are removed and the data (without normalization) is run again, revealing an improved *m* score. SD, standard deviation.

### Technical description of the manifold overlap score for artifact detection

Consider the k-nearest neighborhood of the n^th^ cell in the dataset, which has a baseline condition we define as *x*_*b*_ and a non-baseline condition (eg. IL-7 stimulation) we define as *x*_*i*_ (the i^th^ non-baseline condition). We let *α*_*n*_ be a function that takes in the aforementioned conditions as input (here *x*_*i*_ and x_*b*_), and outputs the count of the cells belonging to a condition *x*_*l*_, divided by the number of cells belonging to both *x*_*i*_ and *x*_*b*_. We define these counts with the subscript n as they pertain to the k-nearest neighborhood of the n^t^^h^ cell.

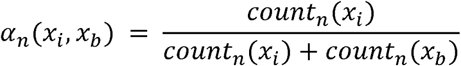

We then let vector *α* (*x*_*i*_, *x*_*b*_) contain all values outputted by *α*_*n*_ (*x*_*i*_, *x*_*b*_) for all *n* cells within the dataset. A different *α* (*x*_*i*_, *x*_*b*_) is calculated for each non-baseline *x*_*i*_ condition in the dataset (as it compares to the user-designated baseline condition *x*_*b*_).

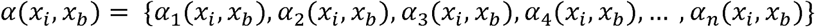

Using *α* (*x*_*i*_, *x*_*b*_) for a given non-baseline condition, we sought to quantify the “overlap” of data between conditions for the expression markers not expected to change, and further test efficacy of normalization and scaling procedures in terms of this overlap. We used the Bodenmiller/Zunder/Finck dataset, and we assessed the *α* of cells treated with GM-CSF. Given that we performed KNN on surface markers that are assumed not to change with a 30-minute *ex vivo* perturbation, we expected the average of each *α* (*x*_*i*_, *x*_*b*_) to be near 0.5 if there were minimal technical artifacts. As an evaluation metric, we calculated the standard deviation of the *α* distributions with each normalization/scaling method, and visually displayed this output as a histogram.

Of note, the software allows the user to determine which markers are not expected to change. The analysis that follows relies on the assumption that any changes in manifold overlap is therefore the result of experimental artifact and not biology. It may of course be found in the future immunology studies that the manifold overlap change we observe here could in fact be biology-related, so the user cannot rule this out.

Our data quality reference was the standard deviation of the binomial distribution with k trials and a probability of 0.5, defined here as *B(k, 0.5).* Our final score was the quotient of the standard deviation of *α* (*s*_1_, *s*_2_) from the subsampled file and *α* (*x*_*i*_, *x*_*b*_) from the baseline condition and “stimulated” condition files. We call this value the manifold overlap score, or *m*.

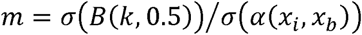

### Performance of manifold overlap score with real and synthetic data

We found that between baseline and GM-CSF conditions, the manifold overlap distribution was substantially different than that of the split-file control (Figure 2B). Per-marker quantile normalization substantially improved this metric minimizing the standard deviation of *α* (*x*_*l*_, *x*_*b*_), and therefore maximizing *m* (Figure 2C). Additionally, identifying problematic markers that went into KNN generation and removing them also improved the *m* score, even without quantile normalization (Figure 2D). Note that it is up to the user to decide whether the problematic markers are of importance to the biological question in mind. If they are, then either normalization or a review of the experimental protocols is recommended.

To determine how sensitive this KNN metric is in detecting marker shifts, we synthetically altered the split-file control showed in Figure 2A. Specifically, we iteratively increased levels of CD4 by half of a standard deviation across all cells in one of the artificial “conditions” of the split-file control. With each iteration, we found the KNN m score and visualized *α* (*x*_*l*_, *x*_*b*_) described above, which we call KNN manifold overlap (Figure S3). We found that a shift of half or a standard deviation decreases *m* from 0.98 to 0.37. This suggests that the m score metric is sensitive enough to detect artifact-driven alterations in a single marker.

We further visualized this synthetic data analysis on t-SNE maps (Figure S4). We show that coloring the tSNE map by CD4 produces a pattern that matches the KNN manifold overlap increasingly with each incremental CD4 shift. We therefore plotted CD4 levels by per-KNN manifold overlap. Of interest, these shifts increase the correlation between the per-cell KNN manifold overlap and CD4 expression levels. This suggests that this type of correlation analysis could be used to efficiently identify markers responsible for a low manifold overlap score, in the event that marker shift has occurred.

Taken together, we recommend the user doing manifold overlap analysis between files in the initial quality control phases of data analysis. The Sconify package allows for this accordingly. Of note, while normalization methods can be of help in certain situations, one cannot possibly rely on a normalization method to “fix” data that has lots of technical artifacts, so special care must always go into the experimental design and data acquisition.

### Visualizing IL-7 responsiveness along the B cell developmental trajectory

The aforementioned Bendall/Davis/Amir dataset was from a study on B cell development using mass cytometry and a specially designed algorithm called Wanderlust to infer a developmental trajectory in static samples of B cell precursors manually gated from healthy human bone marrow (31). This previous study found and focused on a rare population effectively defined by responsiveness to IL-7 through pSTAT5. However, this population’s responsiveness to IL-7 in comparison to other populations could only be interrogated by manual gating or clustering untreated and treated samples.

To directly visualize the location of this subset was within the B cell developmental trajectory, we performed KNN-based marker expression comparisons on these untreated and IL-7 treated samples. We first visualized the data with t-SNE to determine where this population was in relation to related cell surface markers along wanderlust values. The pSTAT5 responsive subset is visible by looking at the t-SNE map of pSTAT5 of the IL7 treated cells only (Figure 3A). This provided “ground truth” to confirm KNN finds this population in its comparisons. The KNN comparison of pSTAT5 between untreated and IL-7 produced a substantially cleaner t-SNE map, colored by pSTAT5 expression differences thresholded by a q value of 0.05 from a Mann-Whitney U test (Figure 3A). Accordingly, this comparison can be added to the Wanderlust Ribbon diagrams used in the original paper. However, we visualize the output as a heatmap in order to maximize the information conveyed (Figure 3B). Wanderlust values are the algorithmically-derived ordering of cells along a virtual developmental trajectory. Our visualization shows that the pSTAT5 responsive population coincides with the initial increase in the expression of specialized polymerase TdT, suggesting high IL7 responsiveness occurs at the beginning of antibody gene rearrangement in B cell precursors (41). Furthermore, this heatmap revealed that the pSTAT5 increase is also coordinated with an increase in CD179a, CD38 along with TdT, and decrease is coordinated with an increase in CD10 along with CD24. Thus, two points of coordinated change in surface marker levels are connected by a small but distinct IL-7 responsive population. We found this architecture to be consistent across four healthy human donors (Figure S5). While we chose a heatmap for visualization here, the KNN-derived change in pSTAT5 added to the expression matrix allows for the user to adapt to any desired visualization without relying on any extra binning steps. Finally, we note that since this B cell development paper came out, there are now a large number of trajectory detection algorithms for multiple-condition single cell data, especially developmental datasets, with which a KNN-based visual approach can be adapted (42).

**Figure 3:**
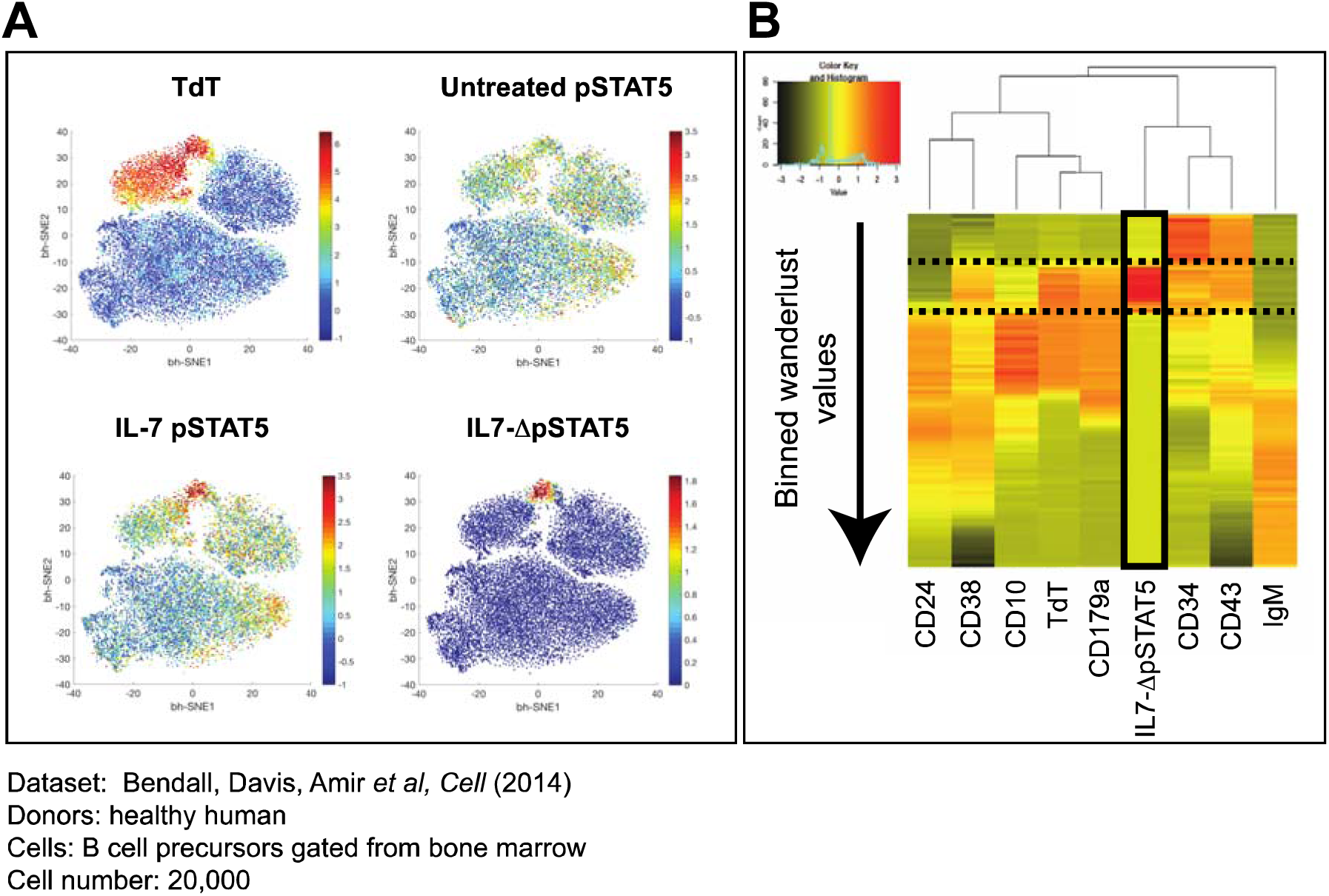
Proof of concept for KNN comparisons. (A) t-SNE map of B cell precursor data. pSTAT5 expression in untreated and IL-7 treated cells reveals ground truth. SCONE produces clean KNN-based change value for an IL-7 responsive subset (B) Heatmap of ordered wanderlust values and KNN-based IL7-pSTAT5 change (IL7-ΔpSTAT5). Dotted lines demarcate the region of increased IL7-ΔpSTAT5, in turn revealing coordination points of marker expression change.

### Runtime analysis

To determine how SCONE performs as a function of the number of cells being used as input, we performed runtime analysis using a comparison between basal and IL10 treated cells in the Fragidakis dataset. This runtime analysis encompassed the KNN generation and the KNN statistics (Figure S6). We omitted the t-SNE step because it is often performed on subsampled data in order to improve runtime, which is done at the user’s discretion. With the KD-tree based KNN generation from the rflann R package, 100,000 cells could be done in 2.3 minutes. The KNN statistics took substantially longer than the KNN generation, with 100,000 cells being done in approximately 13 minutes. The KNN statistics step increased linearly with the increasing number of cells, with every new 10,000 cells taking approximately 1.3 minutes accordingly.

## Discussion

Upon the acquisition of a new dataset, especially that of a new model system or new panel, researchers need to efficiently understand the properties of the dataset enough both identify artifacts in the quality control steps, and to determine the appropriate analysis tools for the questions at hand. Here, we show that KNN-based analysis and visualizations can provide a large amount of relevant information both about experimental artifacts and about the relevant biology. This information can be visualized easily using dimensionality reduction tools, such as t-SNE maps. The ability of SCONE to objectively and automatically set the free parameter (k) allows the user to address the bias-variance tradeoff early in the analysis.

A KNN-derived manifold overlap score was developed and shown to identify batch effects in the Bodenmiller/Zunder/Finck dataset, and showed that normalization and removal of problematic markers could correct for this (Figure 2). We further showed that this manifold overlap analysis was sensitive to synthetically generated alterations in a single marker (Figure S3, S4). This manifold overlap analysis could therefore be used in newly acquired data identify artifacts, find the source of the artifacts (Figure 2), and check for improvement upon removing these artifacts. Outside of dataset quality and computational normalization metrics, manifold overlap analysis could be used to evaluate many experimental conditions, including new or existing blood preservation systems, barcoding systems, and automation systems.

For instances when there are indeed genuine biology-related differences in cell subset abundance between samples, one could use the same architecture to visualize them, thresholding by a defined number of standard deviations from the binomial distribution given the value k, or adapt a EdgeR-based approach (43), such as the one used in Cydar within the KNN architecture (24). Determining whether nearest neighborhoods based on a defined number of a defined distance are more appropriate for this type of analysis is beyond the scope of this paper. However, the rflann package our software uses allows for both types of nearest neighborhoods to be generated. Of note, if the user chooses to user Cydar-generated hyperspheres rather than KNN for expression change analysis, the user could adapt the visualization strategies from the Sconify software accordingly.

The ideal value of k was determined by finding which size of k most effectively imputed the signaling markers of interest in KNN generated by surface markers (Figure S2). While it is beyond the scope of this paper, imputation within the SCONE software is possible by manipulating the KNN list data structure that can be made independent of SCONE. While CyTOF data does not contain missing values, imputation could be used in marker redundancy analysis (29) by determining how well each marker in a panel can be imputed when one leaves it out.

Using a previously published dataset consisting of B cells purified from healthy human bone marrow, we demarcated the boundaries of IL-7 responsiveness through pSTAT5 in relation to other relevant surface markers. Our visualizations showed that both the increase and decrease of IL-7 responsiveness through pSTAT5 across the developmental trajectory was marked by two distinct coordination points, where surface markers abruptly changed as well. Of note, the IL-7-responsive population was previously elucidated by means of a manual gating analysis based on a combination of TdT (high) and CD24 (low) markers (31). We showed that this population can be precisely located and intuitively visualized by SCONE and t-SNE analysis respectively in a fully automated fashion and with single-cell precision, without having to partition the data with gating or clustering. Given that TdT levels are high throughout the IgH gene rearrangement step of developing B cell precursors (44), our visualizations suggest that cells are responsive to IL-7 through pSTAT5 during the early stages of this process. Taken together, our analysis and visualizations effectively summarize the IL-7 pSTAT5 pathway rewiring previously described (31), occuring concomitantly with changes in the expression of multiple surface markers.

SCONE, like t-SNE, is best utilized for visualizations of lower number of cells (eg. 10^4^ to 10^5^) due to run time and multiple hypothesis correction. However, if one were to run SCONE on millions of cells, one could adapt permutation-based statistics to determine which KNN of a user-defined fold change grouped together in a significant manner.

KNN-based analysis along with dimension reduction plots in the early stages of a data analysis pipeline can provide preliminary information about data quality and marker expression changes prior to choosing the appropriate tools(s) for the scaled up data, provided one avoids selection bias. Furthermore, KNN-based analysis in the final stages of analysis can provide effective summary visualizations to show one’s results on a dimension reduction map. Taken together, our tool can be used as a multi-faceted discovery and visualization tool for high-dimensional single cell data, providing the user the ability to compare marker expression and analyze data quality, along with imputing markers, and estimating density within a single method.

### Author contributions

TJB wrote and implemented all code used in this manuscript along with the Bioconductor package, and wrote the manuscript. GPN edited the manuscript and provided direction and guidance. NS provided detailed direction and guidance with both the project and the code, wrote parts of the manuscript, and edited the manuscript.

## Acknowledgements

We would like to acknowledge Kara Davis, and Sean Bendall, and Gabi Fragiadakis whose data was used for this manuscript, and whose insights guided our analysis. We would also like to acknowledge Alyssa Mike and Julie Yu, whose collaboration on a related problem inspired the SCONE approach.

This work was supported by the US National Institutes of Health (grants R33CA0183692, U19AI057229, 1U19AI100627, 7500108142, R01CA184968, 1R33 CA183654-01, 1R01GM10983601, 1R21CA183660, 1R01NS08953301, 5UH2AR067676, 1R01CA19665701, R01HL120724, N01-HV-00242 HHSN26820100034C, 41000411217, 5R01AI073724, U54-UCA149145A to G.P.N), US Department of Defense Congressionally Directed Medical Research Programs (grants OC110674, W81XWH-14-1-0180 to G.P.N), Bill and Melinda Gates Foundation (OPP1113682 to G.P.N), Pfizer, Inc (123214 to G.P.N), US Food and Drug Administration (grants BAA-15-00121, HHSF223201210194C to G.P.N), Novartis Pharmaceuticals Corp (CMEK162AUS06T to G.P.N), Juno Therapeutics Inc (122401 to G.P.N), California Institute for Regenerative Medicine (grant DR1-01477 to G.P.N), Gilead Sciences, Inc. Research Agreement (to G.P.N), NWCRA Entertainment Industry Foundation (to G.P.N), DiaTools (Grant 259796 to G.P.N) Cancer Biology Training Grant 2T32CA009302-36A1 to T.J.B, and the Rachford and Carlota A. Harris Endowed Chair (G.P.N.).

---

**Figure S1:**
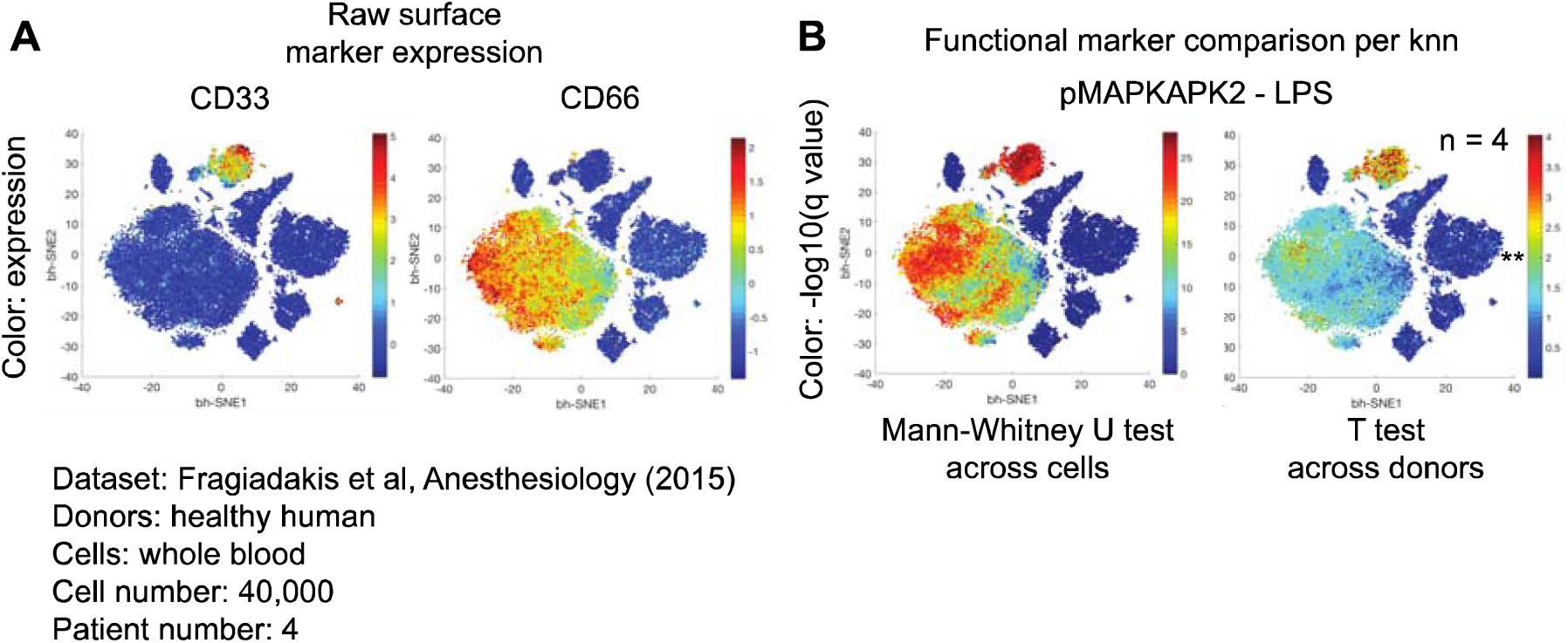
Per-KNN statistical tests can reveal differences both across cells and across donors or replicates. (A) t-SNE maps colored by surface marker expression. (B) Per-cell Mann-Whitney U test or per-donor t test (n = 4) median values of pMAPKAPK2 expression between untreated and LPS-treated cells. ^**^, q < 0.01.

**Figure S2:**
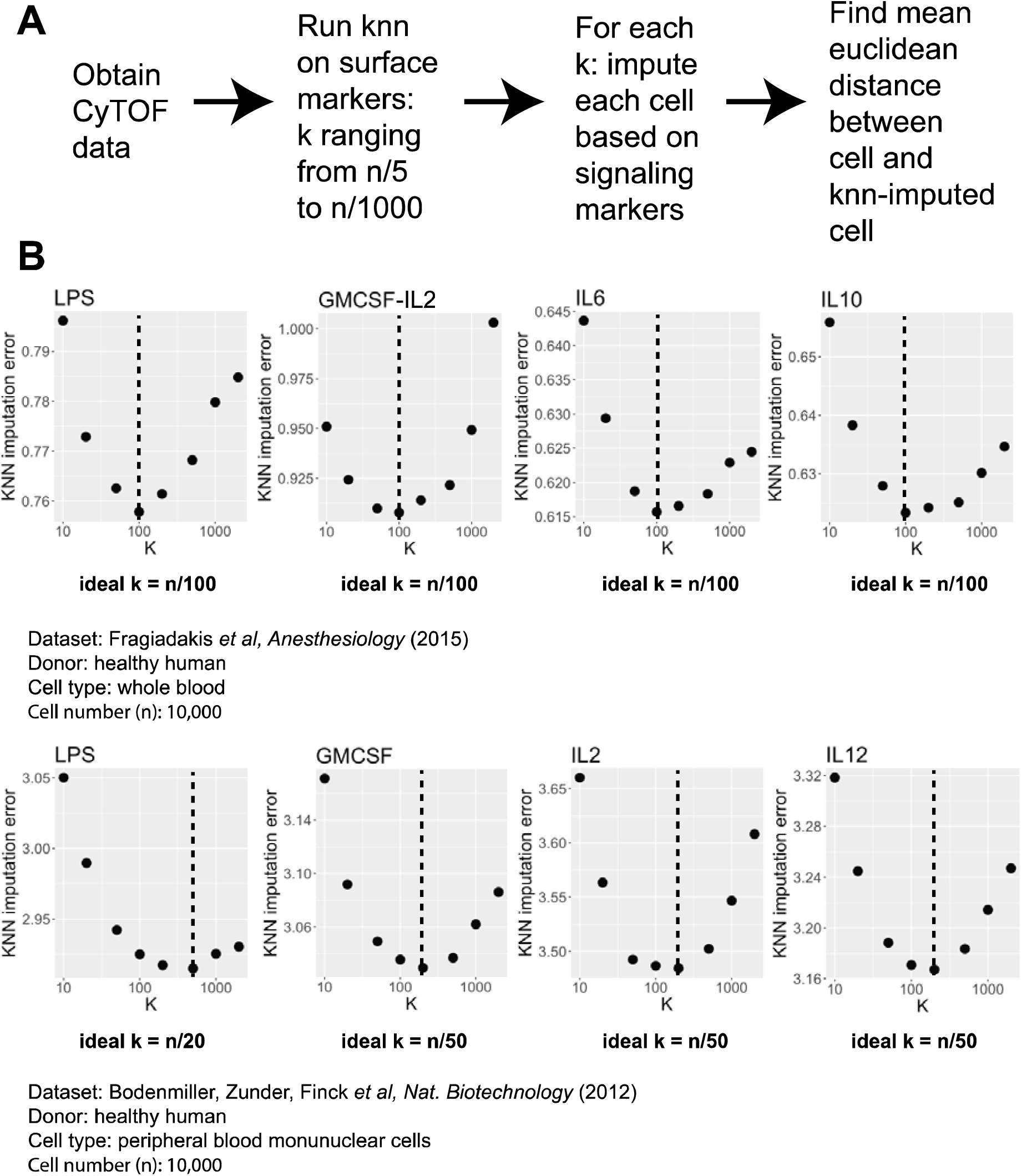
Computational strategy for selection of the number of nearest neighbors (A) Workflow for using various user-selected values of k to impute signaling marker levels from surface markers, and minimizing the error between imputed “cell” and actual cell. (B) Parabolic relationship between selection of (k) and the imputation error in healthy human PBMCs untreated or treated with various immunomodulatory agents in two datasets. Black dashed lines indicate with the minimum imputation error, and therefore the selected value of k. n, number of cells used as input.

**Figure S3:**
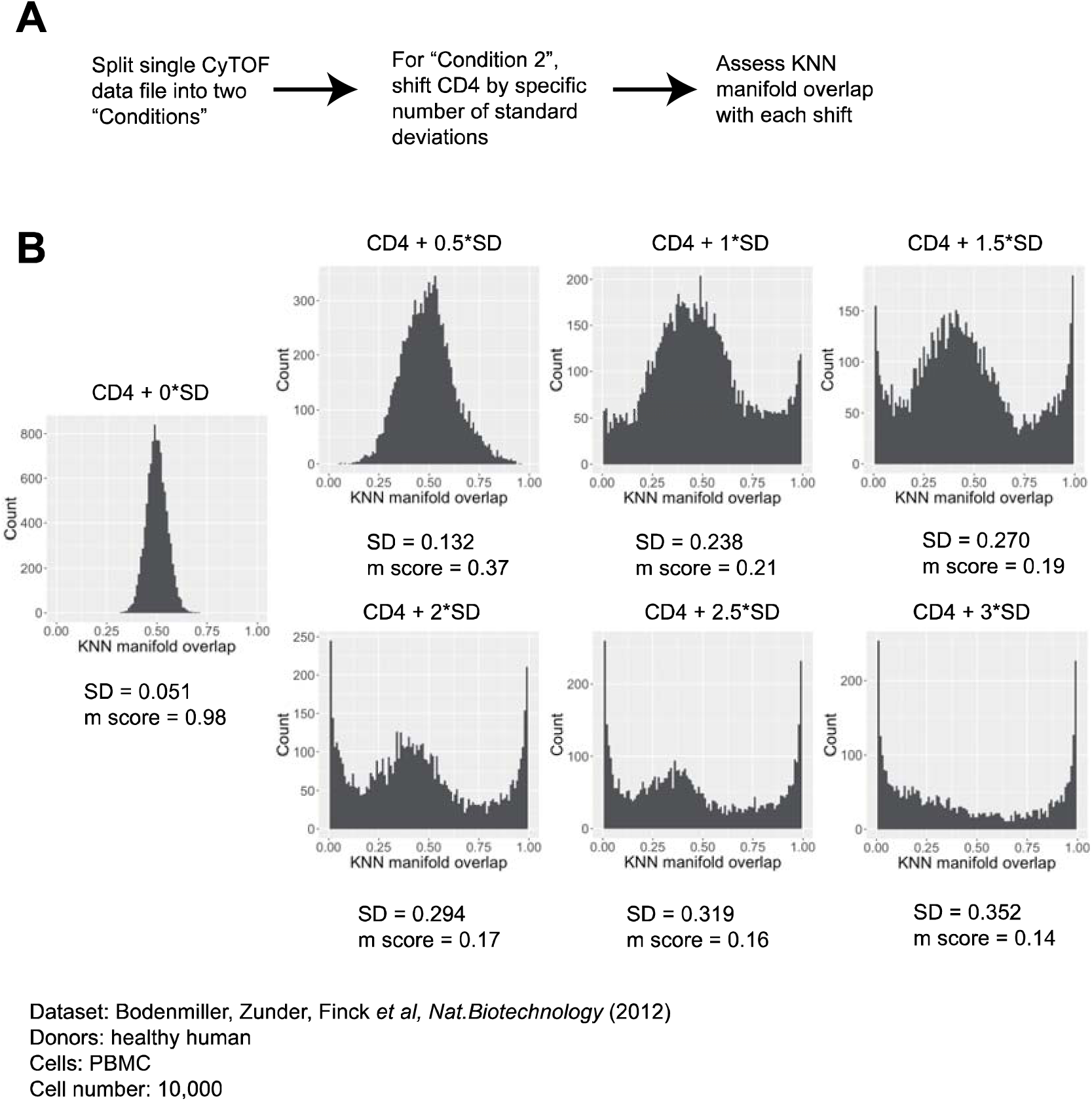
Manifold overlap analysis can detect changes in a single marker. (A) Workflow describing synthetic increase of the CD4 marker by increments of half of its standard deviation. (B) Manifold overlap analysis of the split-file control (see Figure S2A), with increases in CD4 marker levels across one of the two file sub-samples. Increases between 0.5 and 3 standard deviations are tested, with distributions, standard deviations (SD), and *m* score plotted.

**Figure S4:**
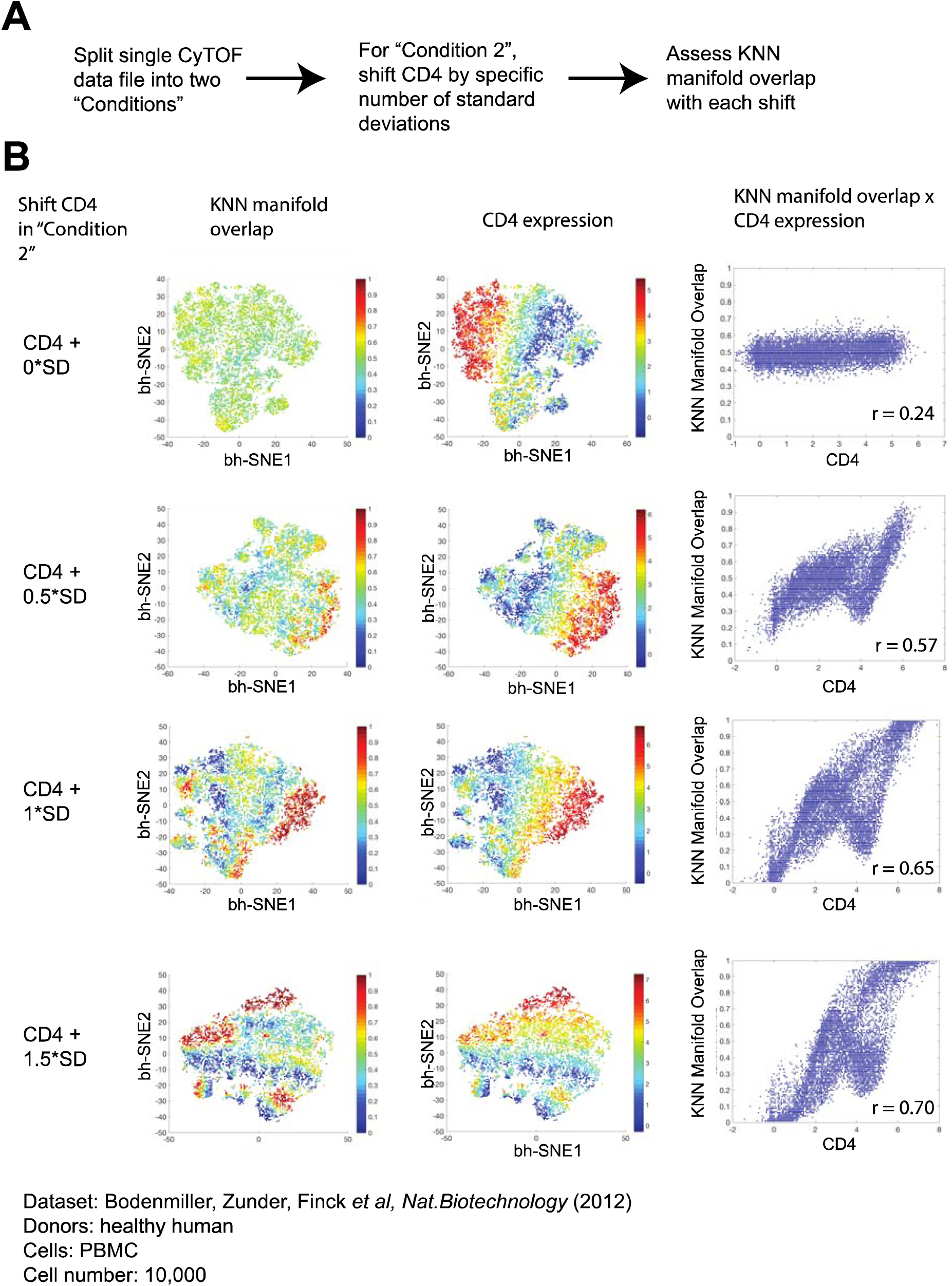
Visualizaiton of manifold overlap analysis from Figure S3 with t-SNE and biaxial plots. (A) Workflow describing synthetic increase of the CD4 marker by increments of half of its standard deviation. (B) (left) tSNE plots colored by CD4 and KNN manifold overlap show that these two parameters are co-expressed in the same regions. (right) Biaxial plots of KNN manifold overlap and CD4 expression reveal a correlation upon synthetic CD4 marker shifting.

**Figure S5:**
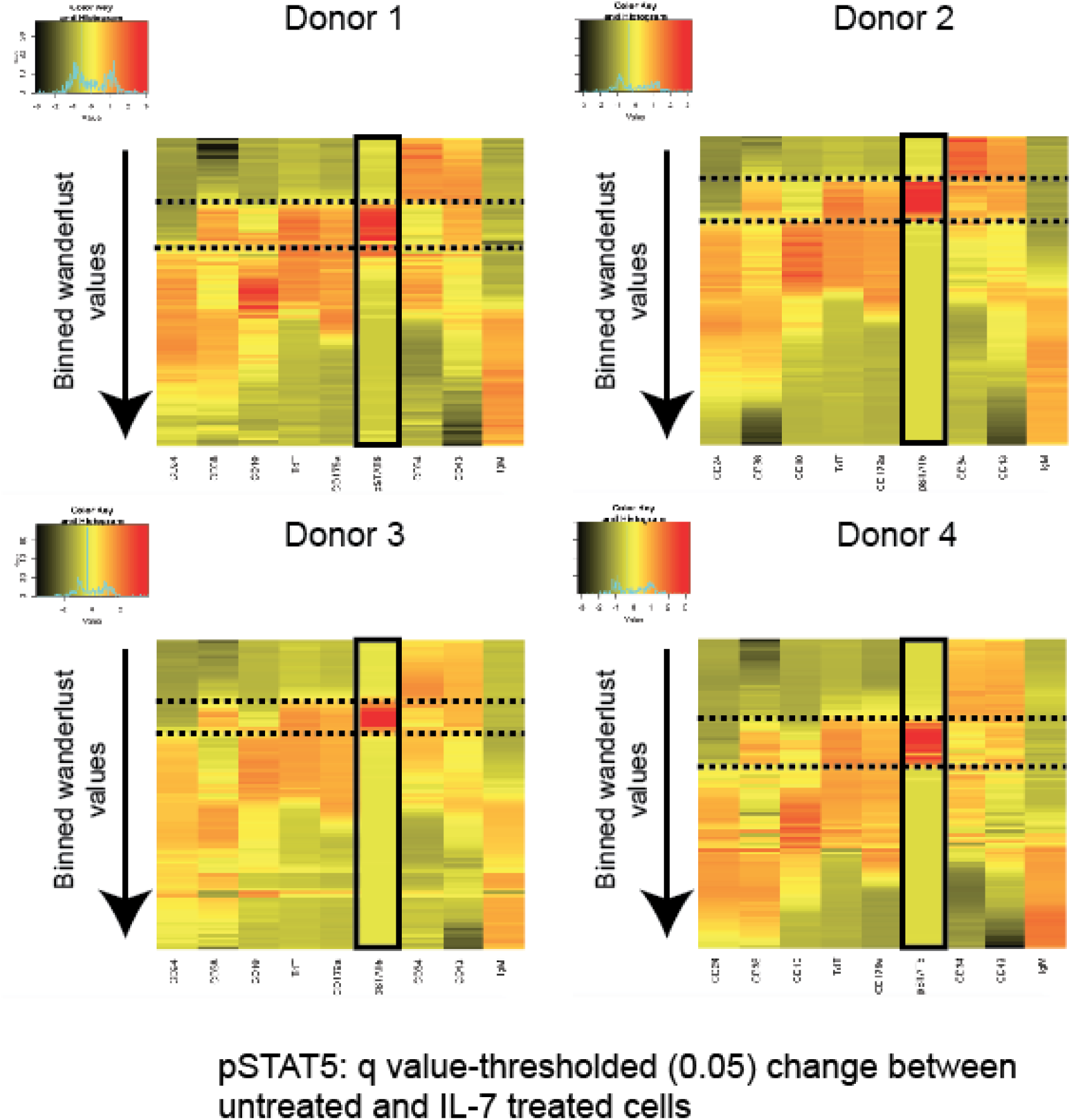
IL7 responsive population through pSTAT5 is consistent across four healthy human donors. Heatmaps for each donor, named donor 1-4, are shown. Wanderlust values are binned, and each bin contains the mean value of the given marker of interest for the cells in that bin. Black box indicates the change in pSTAT5 between untreated and IL7-treated cells for each cell’s given k-nearest neighborhood. Dashed lines indicate “coordination points,” where many surface markers were observed to change simultaneously.

**Figure S6:**
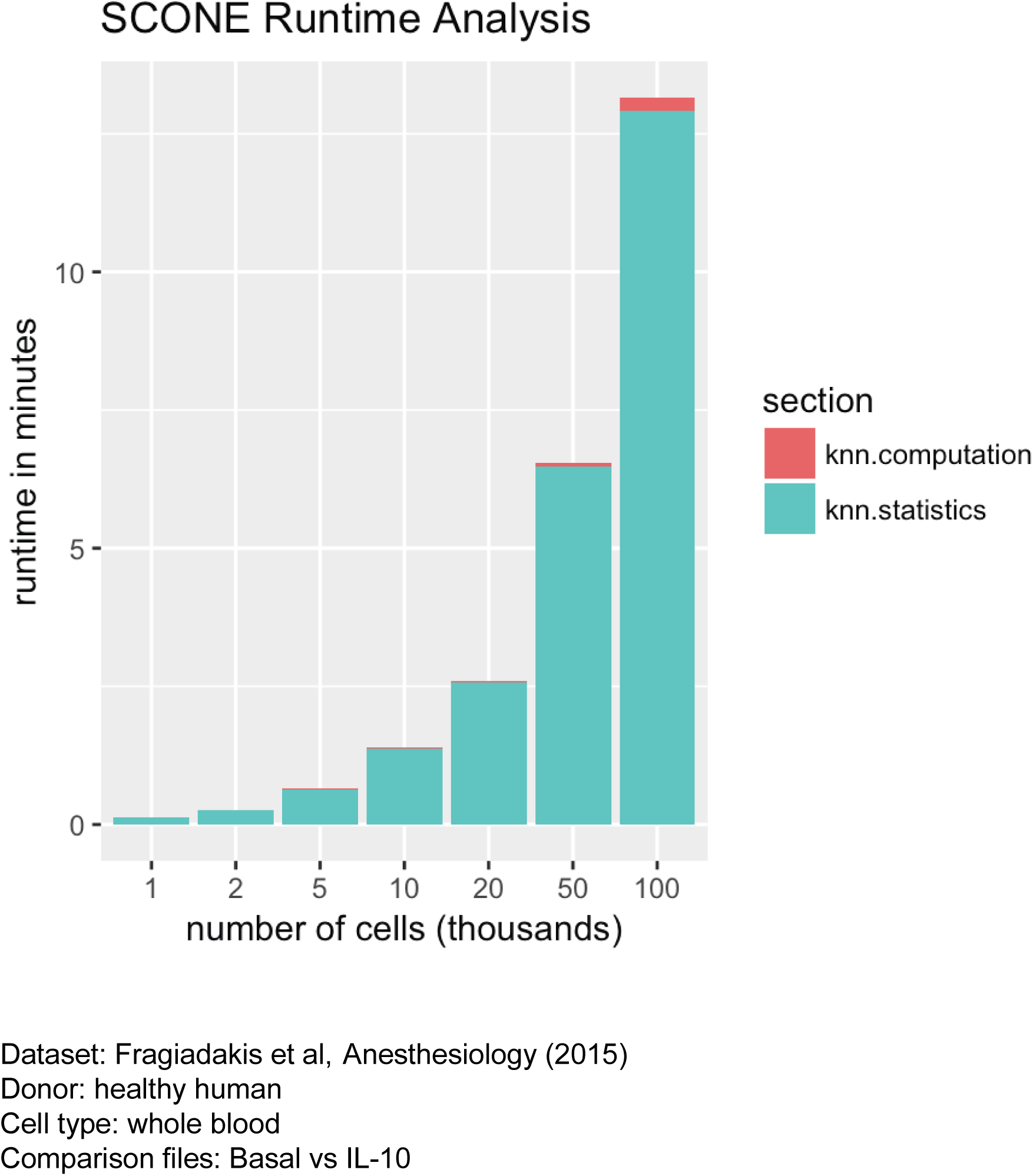
Runtime analysis of KNN computation and KNN statistics for a basal versus IL10 treatment file, across a range of cell numbers between 10^3^ and 10^5^. Total time in minutes includes finding KNN, performing statistical comparisons (Mann-Whitney U test, and q value thresholded change, density, and differential abundance) within these nearest neighborhoods. t-SNE is omitted because it is often done on sub-sampled populations.

